# Atg8 is essential specifically for an autophagy-independent function in apicoplast biogenesis in blood-stage malaria parasites

**DOI:** 10.1101/195578

**Authors:** Marta Walczak, Suresh M. Ganesan, Jacquin C. Niles, Ellen Yeh

**Affiliations:** Department of Biochemistry; Pathology; Microbiology and Immunology, Stanford Medical School; Department of Biological Engineering, Massachusetts Institute of Technology; Chan Zuckerberg Biohub, San Francisco, CA 94158

## Abstract

*Plasmodium* parasites and related pathogens contain an essential non-photosynthetic plastid organelle, the apicoplast, derived from secondary endosymbiosis. Intriguingly, a highly conserved eukaryotic protein, autophagy-related protein 8 (Atg8), has an autophagy-independent function in the apicoplast. Little is known about the novel apicoplast function of Atg8 and its importance in blood-stage *P. falciparum*. Using a *P. falciparum* strain in which Atg8 expression was conditionally regulated, we showed that *Pf*Atg8 is essential for parasite replication. Significantly, growth inhibition caused by the loss of *Pf*Atg8 was reversed by addition of isopentenyl pyrophosphate (IPP), which was previously shown to rescue apicoplast defects in *P. falciparum*. Parasites deficient in *Pf*Atg8, but growth rescued by IPP, had lost their apicoplast. We designed a suite of functional assays, including a new fluorescence *in situ* hybridization (FISH) method for detection of the low-copy apicoplast genome, to interrogate specific steps in apicoplast biogenesis and detect apicoplast defects which preceded the block in parasite replication. Though protein import and membrane expansion of the apicoplast were unaffected, the apicoplast was not inherited by daughter parasites. Our findings demonstrate that, though multiple autophagy-dependent and independent functions have been proposed for *Pf*Atg8, only its role in apicoplast biogenesis is essential. We propose that *Pf*Atg8 is required for fission or segregation of the apicoplast during parasite replication.

## Importance

*Plasmodium* parasites, which cause malaria, and related apicomplexan parasites are important human and veterinary pathogens. They are evolutionarily distant from traditional model organisms and possess a unique plastid organelle, the apicoplast, acquired by an unusual eukaryote-eukaryote endosymbiosis which established novel protein/lipid import and organelle inheritance pathways in the parasite cell. Though the apicoplast is essential for parasite survival in all stages of its life cycle, little is known about these novel biogenesis pathways. We show that malaria parasites have adapted a highly conserved protein required for macroautophagy in yeast and mammals to function specifically in apicoplast inheritance. Our finding elucidates a novel mechanism of organelle biogenesis, essential for pathogenesis, in this divergent branch of pathogenic eukaryotes.

*Plasmodium* (causative agent of malaria) and other apicomplexan parasites are important human and veterinary pathogens. In addition to their biomedical significance, these protozoa represent a branch of the eukaryotic tree distinct from well-studied model organisms that are the textbook examples of eukaryotic biology. As such, parasite biology often reveals startling differences that both highlight the diversity of eukaryotic cell biology and can potentially be leveraged for therapeutic development. A prime example of this unique biology is the non-photosynthetic plastid organelle, the apicoplast. It was acquired by an unusual secondary eukaryote-eukaryote endosymbiosis, in which an alga was engulfed by another eukaryote forming a new secondary plastid in the host (1). Although the apicoplast has lost photosynthetic function, it contains several metabolic pathways and is essential for parasite survival during human infection (2, 3). Despite its importance to pathogenesis, little is known about how the apicoplast coordinates its biogenesis with parasite replication.

*A priori* this unique apicomplexan organelle should have little to do with a highly conserved eukaryotic protein, autophagy-related protein 8 (Atg8). In model organisms, Atg8 plays a central role in autophagy, a conserved eukaryotic pathway for the degradation of cytoplasmic components. During autophagy, cytoplasmic cargo is sequestered in a double-membrane autophagosome which fuses with the lysosome. The ubiquitin-like Atg8 is covalently attached to phosphatidylethanolamine (PE) on the inner and outer membranes of the autophagosome (4). On the autophagosome membrane, it is required for cargo selection, *de novo* formation of the autophagosome and lysosomal fusion, and is the key marker used to identify autophagosomes (5). In fact, blood-stage *Plasmodium* parasites have been reported to accumulate Atg8^+^ vesicles that may represent autophagosomes upon amino acid starvation (6, 7), while Atg8^+^ autophagosome-like structures in liver-stage parasites are required for the turnover of invasion organelles (8).

Yet *Plasmodium* Atg8 clearly has a novel function in the apicoplast, distinct from its role in autophagy. Numerous groups independently showed that Atg8 localizes to the apicoplast in blood-and liver-stage *Plasmodium* as well as the related parasite, *Toxoplasma gondii* (6, 7, 9– 12). Apicoplast localization occurs throughout the parasite replication cycle and is independent of autophagy inducers and inhibitors (7, 9, 13). This function is likely important since the apicoplast is essential for parasite replication during host infection. Indeed, while yeast and mammalian Atg8 homologs are non-essential under nutrient-replete conditions (14, 15), knockdown of Atg8 in *T. gondii* leads to a block in parasite replication with defects in apicoplast biogenesis (16). Consistent with an essential function in *Plasmodium*, Atg7, a component of the Atg8 conjugation system, is essential in blood-stage *P. falciparum* (17), while Atg8 overexpression in liver-stage *P. berghei* results in non-viable parasites with apicoplast defects (8).

Key questions remain: Is Atg8 required for apicoplast biogenesis in the symptomatic blood stage of *Plasmodium falciparum*? It seems likely given the essentiality of *Pf*Atg7 and the phenotypes observed in liver-stage *P. berghei* and *T. gondii* but has not been demonstrated. What is Atg8’s function in apicoplast biogenesis? The abnormal proliferation of apicoplast membranes observed in liver-stage *P. berghei* overexpressing Atg8 was attributed to its role in membrane expansion (8). Meanwhile the association of Atg8 with vesicles containing apicoplast proteins in blood-stage *P. falciparum* suggested a role in vesicle-mediated protein import into the apicoplast (6, 7). Alternatively, *Tg*Atg8 was proposed to mediate the interaction of the apicoplast with the centrosome (16). Since multiple autophagy-dependent and independent Atg8 functions have been proposed, does *Pf*Atg8 have other functions in blood stage essential for parasite replication? For example, Atg8 may have a role in vesicle trafficking to the food vacuole, the lysosomal compartment for host hemoglobin digestion, which is essential for growth in red blood cells (6, 18–20). Atg8’s apicoplast function may be particularly challenging to unravel if other Atg8 functions are also essential.

To answer these questions, we generated a *P. falciparum* strain in which Atg8 expression was conditionally regulated. We assessed parasite replication and apicoplast defects upon Atg8 knockdown, taking advantage of a novel apicoplast chemical rescue only available in blood-stage *P. falciparum*. Not only is *Pf*Atg8 essential for blood-stage *Plasmodium* replication, its only essential function is in apicoplast biogenesis, where it is required for apicoplast inheritance.

## Results

### Atg8 is essential for blood-stage *Plasmodium* replication and apicoplast function

To determine whether *Pf*Atg8 is essential, we generated a conditional expression strain in which the endogenous Atg8 locus was modified with a *C*-terminal myc tag and 3’ UTR tetR-DOZI-binding aptamer sequences for regulated expression (Figure S1). As expected, Atg8 expression was induced in the presence of anhydrotetracycline (aTC) which disrupts the tetR-DOZI repressor-aptamer interaction (Atg8+ condition; Figure 1A and 1C) (21, 22). Though Atg8 was detectable by antibodies against full-length protein, it was not detectable by myc antibodies (Figure S1), suggesting that the *C*-terminus of Atg8 was cleaved. Removal of aTC at the beginning of the parasite replication cycle resulted in efficient knockdown with no detectable Atg8 protein within the same cycle (Figure 1B-C). We monitored the growth of Atg8-deficient parasites and observed a dramatic decrease in parasitemia over 2 or more replication cycles compared to control Atg8+ cultures (Figure 1D). These results show that *Pf*Atg8 is essential for parasite replication in blood-stage *P. falciparum.*

**Figure 1.**
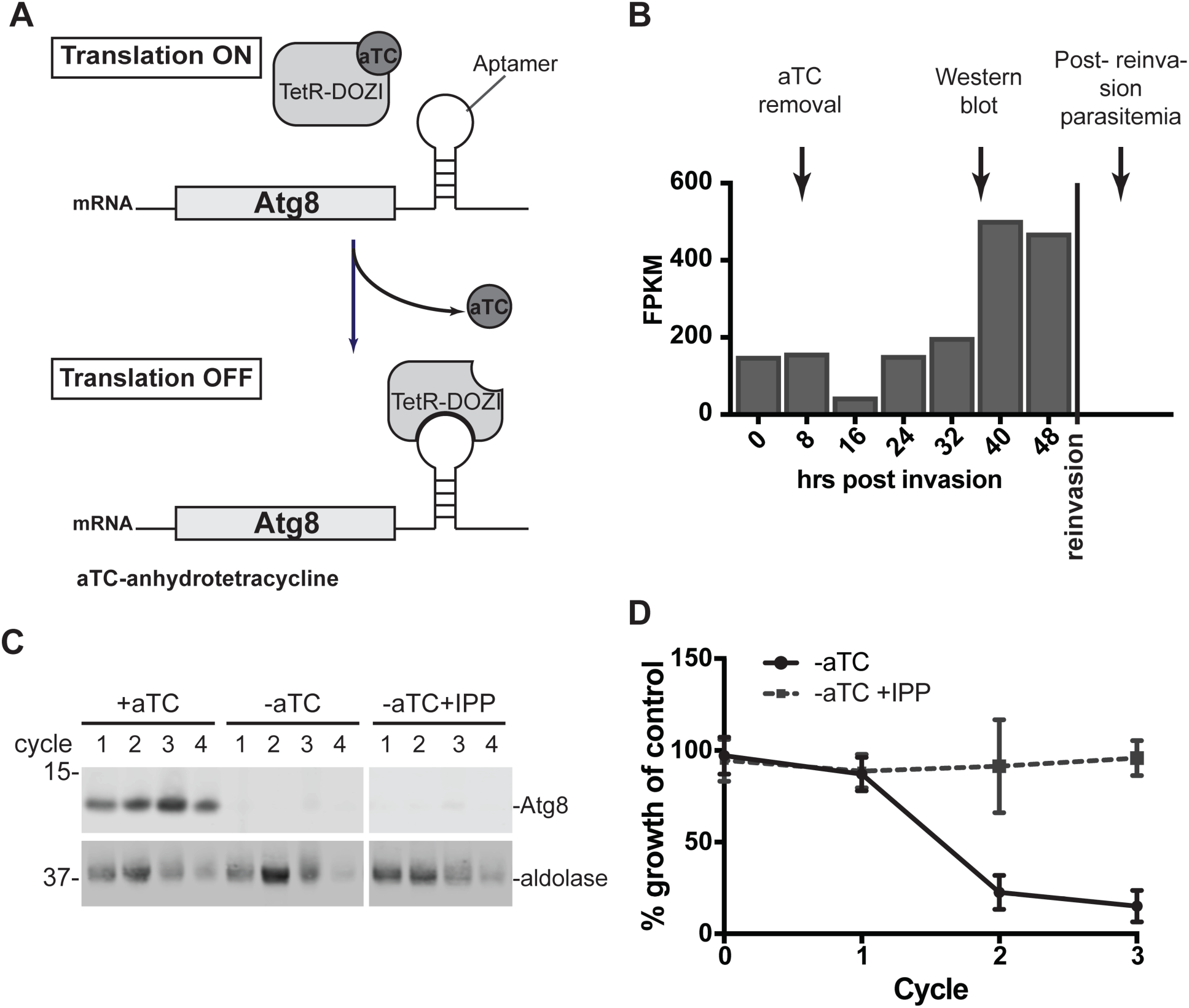
Atg8 is essential for parasite replication and apicoplast function. (A)Regulation of Atg8 expression by anhydrotetracycline (aTC)-dependent binding of TetR-DOZI repressor. (B) Timing of aTC removal and sample collection during a single replication cycle overlaid with Atg8 expression profile (71). (C) Western blot showing Atg8 knock down in the presence or absence of IPP. Equal parasite numbers were loaded per lane. (D) Parasitemia of ACP_L_-GFP expressing cultures grown for 4 cycles under the indicated conditions, normalized to culture grown in the presence of aTC, i.e. expressing Atg8. Average±SD of 3 biological replicates is shown.

The growth inhibition observed in Atg8-deficient parasites may specifically be due to its function in the apicoplast or a result of other functions. To distinguish between essential apicoplast and non-apicoplast Atg8 functions, we determined the growth of Atg8-deficient parasites in media supplemented with isopentenyl pyrophosphate (IPP). We previously showed that IPP is the only essential product of the apicoplast in blood-stage *Plasmodium*. As such, any disruption of the apicoplast, including complete loss of the organelle, can be rescued by the addition of IPP (23). IPP fully rescued the growth defect of Atg8-deficient parasites, demonstrating that the only essential function of Atg8 is specific to the apicoplast (Figure 1D). *Pf*Atg8 may have other functions in blood-stage *Plasmodium* that are not essential but are important for parasite growth fitness, which was not assessed in this study.

### Atg8 depletion leads to apicoplast loss

Each parasite contains a single apicoplast which must be replicated and inherited during cell division. To determine whether *Pf*Atg8 is required for apicoplast biogenesis during parasite replication, we assessed the presence of the apicoplast in Atg8-deficient, IPP-rescued parasites after at least 2 replication cycles when the effects of Atg8 deficiency would be apparent (23). In the first assay, we measured the copy number of the apicoplast genome compared to the nuclear genome and detected a 10-fold decrease in the apicoplast:nuclear genome ratio (Figure 2A). In a second assay, we determined the localization of an apicoplast-targeted GFP (ACP_L_-GFP). In schizont-stage parasites expressing Atg8, ACP_L_-GFP localized to tubular structures which resemble the distinctive branched apicoplast in this stage. In contrast, in Atg8-deficient, IPP-rescued parasites ACP_L_-GFP mislocalized to cytosolic puncta, similar to what has previously been observed in parasites in which apicoplast loss has been induced by treatment with apicoplast transcription and translation inhibitors like chloramphenicol (Figure 2B-C) (23, 24). Altogether, these results indicate that the apicoplast is lost in Atg8-deficient parasites, likely due to a failure to replicate and inherit new apicoplasts during parasite replication.

**Figure 2.**
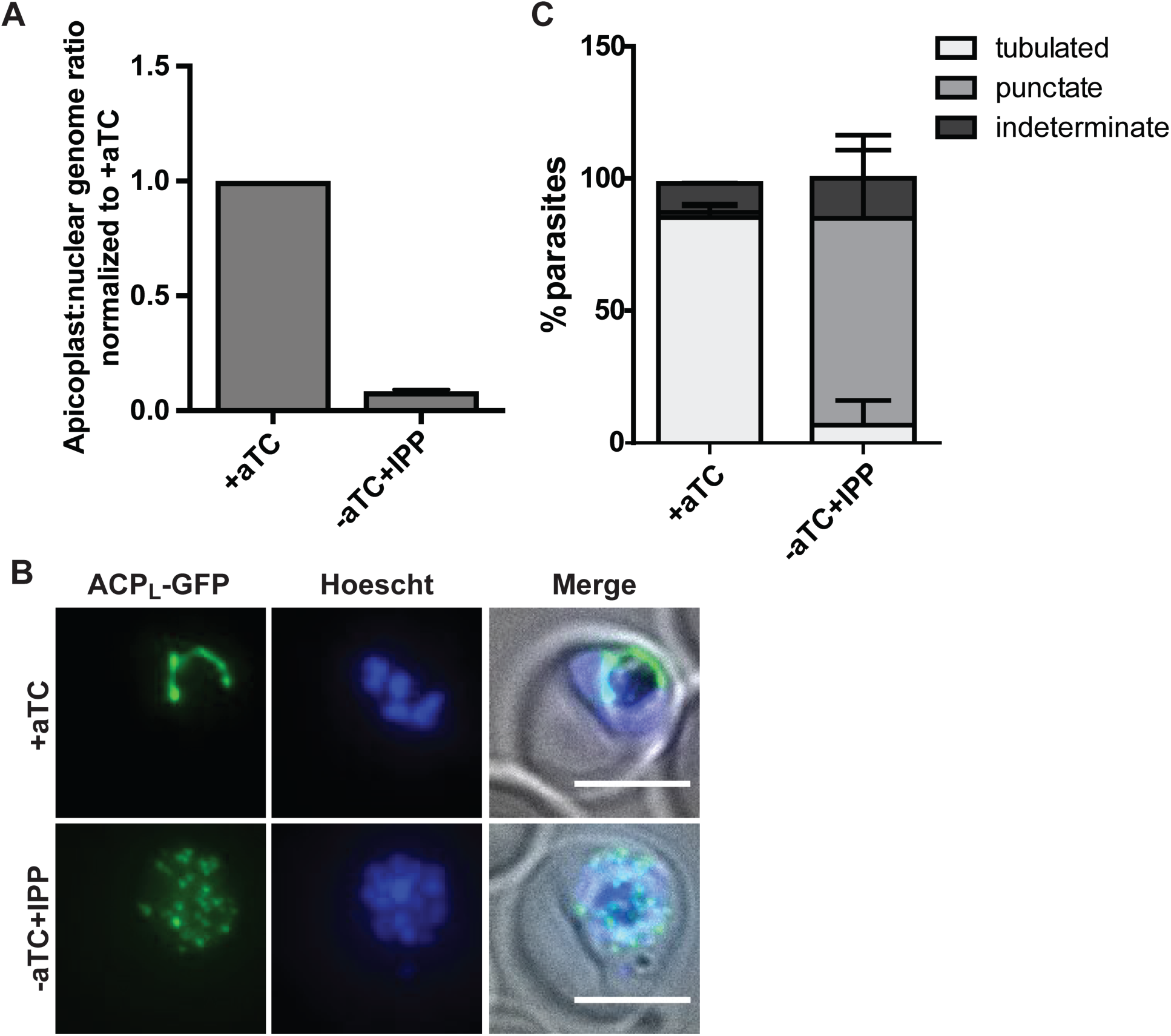
Atg8 depletion leads to apicoplast lost. (A)Apicoplast:nuclear genome ratio in Atg8-deficient, IPP-rescued parasites (grown for 4 cycles without aTC) measured by quantitative PCR. The ratios were normalized to Atg8+ culture (i.e. grown in the presence of aTC). (B) Representative microscopy images showing localization of apicoplast-targeted GFP (ACP_L_-GFP), in schizont-stage Atg8+ or Atg8-deficient/IPP rescued parasites depleted of Atg8 for 2 replication cycles. Scale bar, 5 μm. (C) Quantification of parasites with the indicated apicoplast morphology in Atg8+ or Atg8-deficient/IPP rescued parasites as shown in B. Average±SD of 2 independent experiments is shown.

### Atg8 depletion does not affect protein and lipid import to the apicoplast

We noted that parasite growth was initially unaffected by Atg8 knockdown but then decreased drastically in the subsequent replication cycle. As seen in Figure 1C-D, despite substantial Atg8 depletion upon aTC removal, Atg8-deficient parasites reinvaded new host cells efficiently achieving similar parasitemia as Atg8+ parasites in cycle 1. However, in the subsequent reinvasion (cycle 2), the parasitemia was 26% of the control. To determine whether Atg8 depletion caused defects in apicoplast biogenesis in cycle 1 that preceded the block in parasite replication in cycle 2, we monitored key events in apicoplast biogenesis in the first cycle of Atg8 knockdown (Figure 3A).

**Figure 3.**
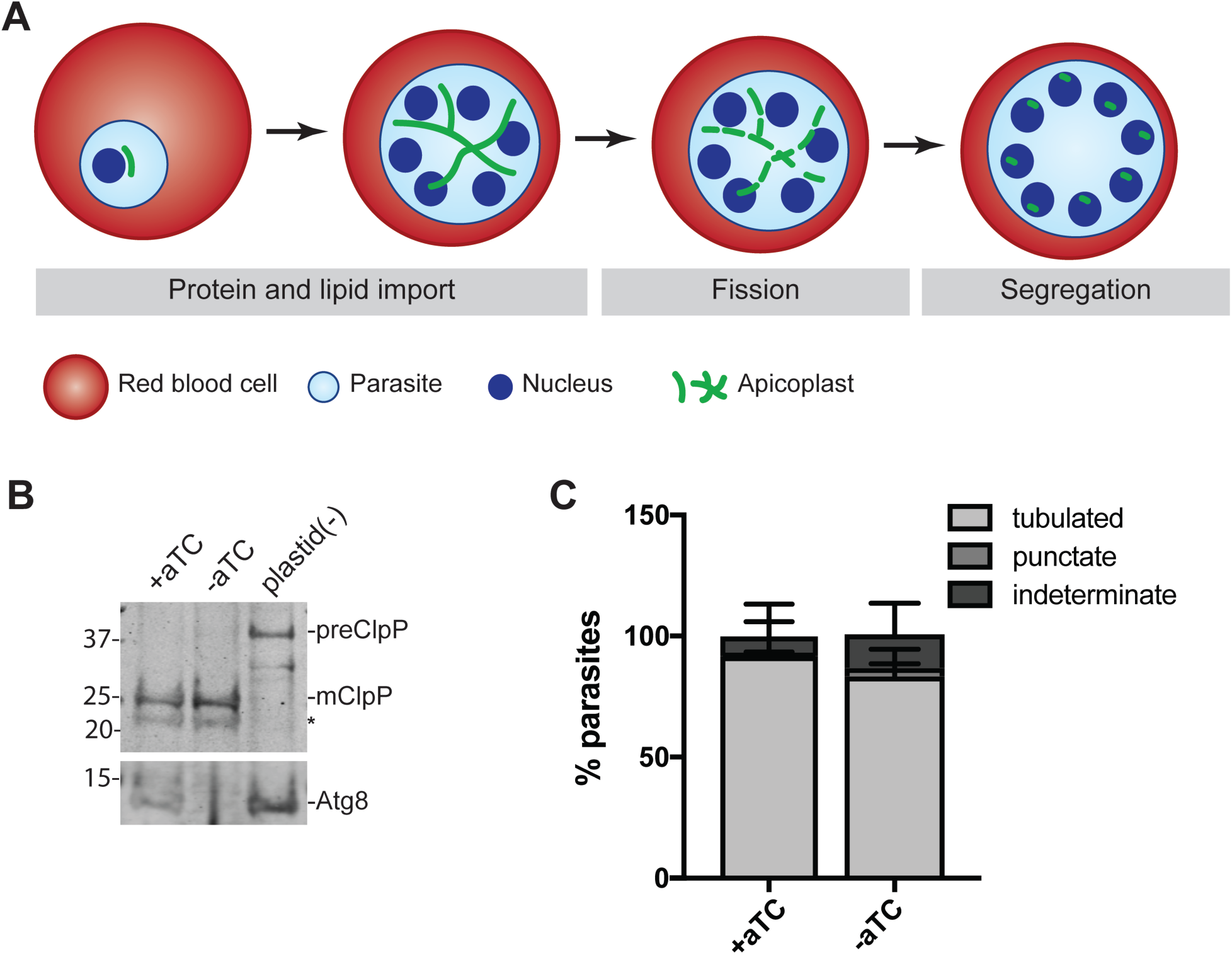
Apicoplast protein import and membrane expansion are not affected in the first cycle of Atg8 knockdown. (A)Time course of molecular events during apicoplast development in blood stage parasites. (B) Processing of a luminal apicoplast protein, ClpP, in Atg8+ or Atg8-deficient parasites approximately 24 hrs post aTC removal. Apicoplast(-) parasites generated by chloramphenicol treatment and IPP rescue over 4 replication cycles (23), which possess only precursor ClpP, are shown for reference. preClpP, full length (precursor) form of ClpP, 43 kDa; mClpP, mature (apicoplast-luminal) ClpP, 25 kDa. The asterisk indicates a non-specific band. Atg8 expression in the corresponding time points is shown for reference. (C) Quantification of parasites with the indicated apicoplast morphology during the first cycle of Atg8 knockdown 32 hrs after aTC removal. Apicoplast was visualized using the luminal apicoplast marker, ACP_L_-GFP. Representative images are shown in Figure S2.

The first distinctive change associated with apicoplast biogenesis is growth and formation of a branched apicoplast (25, 26), which is likely dependent on protein and lipid import. Apicoplast-targeted proteins possess an *N*-terminal transit peptide sequence which targets them to the apicoplast and is removed upon import into the apicoplast (27). To assess apicoplast protein import in Atg8-deficient parasites, we monitored the processing of an imported protein, ClpP, from a 43 kDa full-length protein containing an intact transit peptide (as observed in chloramphenicol-induced apicoplast loss) to a 25 kDa mature form (28). We observed no defect in ClpP processing in trophozoite parasites ∼24 hours after Atg8 knockdown (Figure 3B). Furthermore, apicoplast-targeted ACP_L_-GFP localized to a branched tubular structure similar to those in Atg8+ parasites in schizont parasites ∼32 hours after Atg8 knockdown, indicating that lipid import contributing to this extensive membrane expansion was also unaffected (Figure 3C and S2). Our data suggest that Atg8 expression is not immediately required for apicoplast protein import or membrane expansion.

### Atg8 knockdown results in a late block in apicoplast inheritance

The final events in apicoplast biogenesis are division of the branched apicoplast to form multiple plastids and segregation of a single apicoplast into each forming daughter parasite (merozoite). These events required for organelle inheritance have not been directly observed. Instead we assessed apicoplast inheritance upon Atg8 knockdown by detecting the presence of the apicoplast genome and apicoplast-targeted ACP_L_-GFP in newly reinvaded Atg8-deficient parasites (48 hours after aTC removal and ∼12 hours post-invasion) after the first cycle of Atg8 knockdown. As noted, Atg8-deficient parasites reinvaded to similar parasitemia as Atg8+ parasites in this first reinvasion (Figure 1D).

To detect the single-copy apicoplast genome with single cell resolution we developed a fluorescence *in-situ* hybridization (FISH) protocol using an Oligopaints library of 477 FISH probes covering >60% of the 35 kb genome (apicoplast FISH) (29, 30). As expected, a majority of Atg8+ parasites (85%) had a single fluorescent punctum corresponding to the apicoplast genome (Figure 4A-B and S3). This punctum was absent from negative-control parasites in which apicoplast loss had been induced by chloramphenicol treatment, demonstrating that apicoplast FISH was specific (Figure 4A-B) (23). In contrast to Atg8+ parasites, only 19% of reinvaded Atg8-deficient parasites contained an apicoplast genome after the first cycle of Atg8 knockdown (Figure 4A-B). Since the experiments were performed on a non-clonal population, the small percentage of apicoplast FISH-positive parasites in the Atg8-deficient pool was likely due to incomplete Atg8 knockdown or unmodified wildtype parasites.

**Figure 4.**
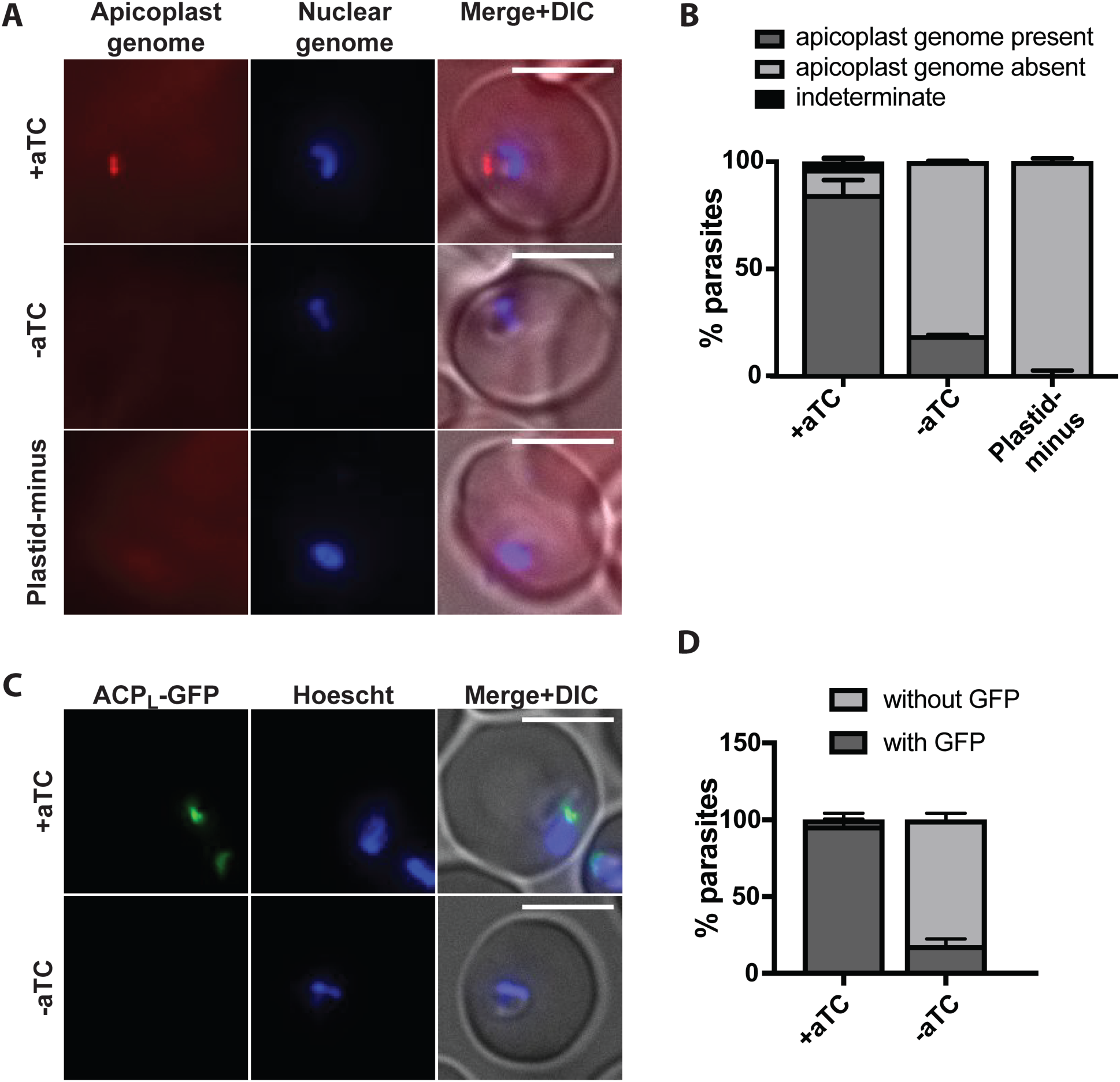
Atg8 knockdown leads to defects in apicoplast inheritance. (A) Representative images of apicoplast FISH detecting the apicoplast genome in ring stage Atg8+, Atg8-deficient, and apicoplast-minus parasites as a negative control for FISH staining. Apicoplast-minus parasites generated by 4 cycles of chloramphenicol treatment and IPP. Scale bar 5 μm. (B) Quantification of parasites with or without apicoplast genome grown under indicated conditions. Average±SD of 2 independent experiments as in (A) is shown. (C) Representative images of Atg8+ or Atg8-deficient parasites (48 hrs post aTC removal) expressing ACP_L_-GFP. Scale bar 5 μm. (D) Quantification of parasites with or without a discrete GFP-labelled structure as shown in (C) grown under indicated conditions. Average±SD of 2 independent experiments is shown.

Similarly, we detected ACP_L_-GFP in individual ring-stage parasites by fluorescence microscopy (Figure 4C-D). Consistent with apicoplast FISH results, 96% Atg8+ parasites had a punctate or elongated ACP_L_-GFP signal, while only 18% Atg8-deficient parasites contained detectable ACP_L_-GFP. The presence of the apicoplast genome and protein in these early ring-stage parasites should reflect inheritance of the organelle, rather than DNA or protein synthesis, since neither genome replication nor GFP expression is active in this stage. Therefore, the decrease in apicoplast FISH and ACP_L_-GFP labelled structures in the progeny of Atg8-deficient parasites suggests that apicoplast inheritance was disrupted.

## Discussion

Our findings demonstrate that *Pf*Atg8 has a novel, essential function in apicoplast biogenesis which is conserved among apicomplexan parasites. *Pf*Atg8, like the Atg8 homolog in *T. gondii*, is essential for parasite replication. The essentiality of apicomplexan Atg8 contrasts with yeast and mammalian Atg8 homologs which are not strictly required for cell growth and proliferation in nutrient-replete conditions (14, 15). Moreover, though Atg8 has been proposed to have diverse functions in *Plasmodium* parasites from starvation-induced autophagy to stage-specific organelle turnover to intracellular vesicle trafficking (6–8), we showed that *only* its role in apicoplast biogenesis is essential for blood-stage *Plasmodium* replication. *Tg*Atg8’s essentiality was also attributed to its apicoplast function since neither autophagosome biogenesis by Atg9 nor proteolysis in the lysosomal compartment is essential in replicating tachyzoites (31– 33). This unique function of *Pf*Atg8 may be leveraged for antimalarial drug development. Since autophagy has important roles in mammalian physiology and development, specificity for disruption of *Pf*Atg8 and its conjugation will be imperative. One strategy may be to identify druggable targets downstream of *Pf*Atg8 that specifically affect apicoplast biogenesis (15, 34), though it is unclear whether direct inhibition of Atg8 function (as opposed to interfering with its expression) will result in the delayed growth inhibition observed in our Atg8 knockdown strain. Finally, we determined essential *Pf*Atg8 functions for blood-stage *P. falciparum* growth using an *in vitro* culture system; it is possible that Atg8 has other essential functions under *in vivo* conditions and/or in other life stages.

*Pf*Atg8 has a novel function in apicoplast biogenesis. Because Atg8 homologs in model eukaryotes have not previously been implicated in biogenesis of mitochondria or primary chloroplasts, this function likely evolved as a result of secondary endosymbiosis in this parasite lineage. The repurposing of a conserved eukaryotic protein for the biogenesis of a secondary plastid is at first surprising. However, Atg8-conjugated membranes of the apicoplast and autophagosomes both have their origins in the endomembrane system. The ER and ER-associated membranes are main membrane source of autophagosomes (35–40). Meanwhile Atg8 is conjugated to the outermost of 4 apicoplast membranes (16), which derives from the host endomembrane during secondary endosymbiosis (1). Indeed, apicoplast biology has numerous tantalizing connections to ER biology. Protein import into the apicoplast requires that nuclear-encoded proteins traffic to the ER *en route* to the apicoplast (27). A translocon related to the ER-associated protein degradation (ERAD) system localizes to the apicoplast and may be involved in protein import, another example of an ER-associated membrane function that has been repurposed for apicoplast function (41–44). Finally, in some free-living protists, the secondary plastid is located within the ER with the outermost membrane of the plastid contiguous with the ER (45, 46). The endomembrane origin of the outer membrane may explain the novel function of Atg8 in apicoplast biogenesis.

What is the function of Atg8 on this outermost membrane? Mammalian and yeast Atg8 homologs have two unique properties that contribute to their diverse autophagy-related and autophagy-independent functions. First, they stimulate membrane tethering, hemifusion, and fusion, important for their role in autophagosome formation (47, 48). Based on this membrane fusion activity, *Pf*Atg8 was proposed to promote membrane expansion of a growing apicoplast and/or fusion of vesicles containing nuclear-encoded proteins with the apicoplast (6–8, 10, 12, 49). However, in Atg8-deficient *P. falciparum*, we did not observe any defect in either membrane expansion (assayed by formation of a branched intermediate) or protein import (assayed by ClpP transit peptide processing) prior to the block in parasite replication. We therefore consider these putative functions less likely.

Second, ubiquitin-like Atg8 proteins are versatile protein scaffolds for membrane complexes, interacting with a variety of effector proteins, including cargo receptors, SNAREs, NSF, Rab GAPs, and microtubules (50–56). In Atg8-deficient parasites, we observed a defect in apicoplast inheritance, resulting in loss of a functional apicoplast in their progeny. We propose that *Pf*Atg8 is required to resolve the branched intermediate into individual apicoplasts (fission) and/or facilitate the distribution of a single apicoplast into each budding daughter parasite (segregation). Are there known Atg8 effectors that provide a model for these functions? To our knowledge, interaction of Atg8 homologs with normal-topology membrane fission machinery, such as dynamins, has not been reported (57). Mammalian Atg8 homologs, LC3 and GABARAP, have been shown to interact directly and indirectly with centrosomal proteins, albeit not conserved in apicomplexans (58–60). In *T. gondii* and another apicomplexan *Sarcocytis neurona,* dividing apicoplasts are associated with centrosomes, which may serve as a counting mechanism to ensure inheritance of a single apicoplast by each daughter parasite (61, 62). Notably, the association is independent of the mitotic spindle and lost upon knockdown of Atg8 in *T. gondii*, suggesting that Atg8 may mediate this interaction with centrosomal proteins (16, 62). Though *Plasmodium* lacks centrioles and instead contains “centrosome-like” structures, apicoplast-bound Atg8 may interact with these structures in *Plasmodium* as well (63, 64). Finally, LC3 and GABARAPs also interact with microtubules and may be required for the transport of autophagosomes and GABA receptor-containing vesicles, respectively (53, 55, 56, 65). By analogy, *Pf*Atg8 may interact with microtubules to position the apicoplast during parasite division. Indeed, *Pf*Atg8 may interact with multiple effectors at the apicoplast membrane, as it does on autophagosomes, to ensure organelle inheritance. Identifying these effectors will be a challenging but critical next step.

Atg8’s function in apicoplast biogenesis is required in different life stages of *Plasmodium* spp and conserved with related apicomplexan parasites. In fact, the apicoplast function of apicomplexan Atg8 is the most consistently observed. Our results in blood-stage *Plasmodium* corroborate findings in liver-stage *Plasmodium* and *T. gondii* tachyzoites that also showed a role in apicoplast biogenesis (8, 10, 12, 16). Even autophagy, which is the “ancestral” function of Atg8, is not clearly preserved in *Plasmodium* parasites. It will be interesting to determine whether Atg8’s role in apicoplast biogenesis is a specific adaptation of apicomplexan parasites or is also found in free-living relatives that possess a secondary plastid of the same origin such as *Chromera*. Overall the evolution of this new protein function for a key endosymbiotic event from an ancient template is intriguing (66).

## Materials and methods

### Culture and transfection conditions

*Plasmodium falciparum* parasites were grown in human erythrocytes (Research Blood Components, Boston, MA/Stanford Blood Center, Stanford, CA) at 2 % hematocrit under 5% O_2_ and 5% CO_2_, at 37° C in RPMI 1640 media supplemented with 5 g/l Albumax II (Gibco), 2 g/l NaHCO_3_ (Fisher), 25 mM HEPES pH 7.4 (Sigma), 0.1 mM hypoxanthine (Sigma) and 50 mg/l gentamicin (Gold Biotechnology) (further referred to as culture medium). For transfections, 50μg plasmid DNA were used per 200 μl packed red blood cells (RBCs), adjusted to 50% hematocrit, and electroporated as previously described (21). Parasites were selected with a combination of 2.5 mg/l blasticidin S (RPI Research Products) and 2.5 nM WR99210 (Atg8 TetR strain) or 2.5 mg/l blasticidin S and 500 μg/ml G418 sulfate (Corning) (ACPL-GFP expressing Atg8-TetR strain) beginning 4 days after transfection.

### Cloning and strain generation

All primers used for this study are listed in Supplementary Table 1. *P. falciparum* NF54^attB^ parasites (kindly provided by David Fidock) engineered to continuously express Cas9 and T7 RNA polymerase (NF54^Cas9+T7 Polymerase^) (67) were used as parental strain for deriving Atg8 conditional knock down parasites.

The construct for inducible Atg8 expression, pSN053-Atg8, was created by cloning left and right homology arms and guide RNA into a pJazz system based vector, pSN053. The pSN053 vector contains 1) a C-terminal myc tag followed by 10x Aptamer array for anhydrotetracycline-dependent regulation of translation, 2) a TetR-DOZI cassette containing Renilla luciferease (RLuc) gene for monitoring transfection, blasticidin resistance gene for selection, and a TetR-DOZI repressor, with PfHrp2 3’ and PfHsp86 5’ regulatory sequences, in head-to-head orientation with the modified gene, and 3) a guide RNA expression cassette with T7 promoter and T7 terminator. The left homology arm was amplified from genomic DNA with primers SMG413+SMG425, and inserted into FseI-AsiSI site in frame with the myc tag. The right homology arm was amplified from genomic DNA with the primers SMG411+SMG412 and inserted into the I-SceI and I-CeuI sites downstream of the TetR-DOZI cassette. The guide RNA was generated by Klenow reaction from oligonucleotides SMG419 and SMG420 and inserted into the AflII site. All ligation steps were performed using Gibson assembly. The resulting plasmid was transfected into the NF54^Cas9^+T7 ^Polymerase^ strain as described above and transformants were selected with 2.5 μg/l blasticidin S and 2.5 nM WR99210. Culture was maintained in 0.5μM anhydrotetracycline (aTC) (Sigma) unless stated otherwise. Transgene integration (5’ junction) was confirmed by PCR using primers SMG454 and SMG493. We were not able to amplify a product on the 3’ junction. This strain is referred to as Atg8-TetR strain.

To introduce a fluorescent apicoplast marker, GFP with an apicoplast targeting leader sequence, ACP_L_-GFP was amplified from pRL2-ACP_L_-GFP using primers mawa059 and mawa060 and cloned into AvrII-SacII restriction sites of a pY110F plasmid using InFusion (Clontech). The plasmid was transfected into Atg8-TetR strain and transformants were selected with 2.5 mg/l blasticidin S and 500 mg/l G418 sulfate. Cultures were maintained in 0.5 μM aTC and 500 mg/l G418 sulfate.

### Atg8 knock down experiments

Ring stage parasites at 5-10% parasitemia were washed twice in the culture medium to remove aTC, resuspended in the culture medium and the hematocrit was adjusted to 2 %. Parasites were divided into 3 cultures grown in the culture medium supplemented with 0.5 μM aTC, without aTC, or without aTC with 200 μM IPP (Isoprenoids) for 4 replication cycles. At schizont stage of each cycle, cultures were diluted 5-fold into fresh culture media with red blood cells at 2 % hematocrit and aTC or IPP was added as required. Aliquots of culture for western blot, quantitative PCR and flow cytometry were collected at ring and/or schizont stage of each cycle, before diluting the cultures.

### Flow cytometry

Parasite cultures or unifected RBCs at 2 % hematocrit were fixed with 1% paraformaldehyde (Electron Microscopy Solutions) in PBS for 4 hours at RT or overnight at 4° C. Nuclei were stained with 50 nM YOYO-1 (Life Technologies) for minimum 1 hour at room temperature. Parasites were analyzed on the BD Accuri C6 flow cytometer. Measurements were done in technical triplicates.

### Western blot

Parasites were lysed with 1 % saponin for 5 min on ice. Parasite pellets were washed twice with ice-cold PBS and resuspended in 20 μl 1x LDS buffer (Life Technologies) per 1 ml culture at 2% hematocrit, 5 % parasitemia. Equal parasite numbers were loaded per lane. After separation on Bis-Tris Novex gels (Invitrogen), proteins were transferred to a nitrocellulose membrane, blocked with a buffer containing 0.1 % casein (Hammarsten, Affymetrix), 0.2x PBS and incubated with the corresponding antibodies diluted in 50% blocking buffer/50% TBST. Primary antibodies were used overnight in 1:1,000 dilution, except anti-aldolase which was used at 1:10,000 and anti-GFP used at 1:20,000. Secondary antibodies were used at 1:10,000 dilution for 1 hour at room temperature. Blots were visualized using Licor double-color detection system and converted to grayscale images for the purpose of this publication. Following antibodies were used: anti-Atg8, Josman LLC (see below); anti-aldolase (Abcam ab207494), anti-ClpP, a gift from W. Houry (28); anti-GFP (Clontech 632381). Fluorophore-conjugated IRDye secondary antibodies were purchased from Fisher (Licor).

### Quantitative PCR

0.5 ml culture were lysed with 1 % saponin and washed twice with PBS. DNA was purified using the DNeasy Blood and Tissue kit (Qiagen). PCR reactions were prepared using LightCycler 480 SYBR Green I Master mix (Roche) according to manufacturer’s instructions and run in triplicates on the Applied Biosystem 7900HT cycler. Primers TufA fwd and TufA rev were used for the apicoplast target, and Cht1 fwd and Cht1 rev for the nuclear target. Cycling conditions were: 95° C-10 min; 35 cycles of 95° C-30 s, 56° C-30 s, 65° C-90 s; 65°C-5 min; melting curve 65°-95° C. Data were analyzed using a delta-delta C_T_ method as previously described (68).

### Fluorescence microscopy

Live or fixed parasites were stained with 2 μg/ml Hoescht 33342 stain for 15 min at room temperature to visualize nuclei. Images were acquired using the Olympus IX70 microscope equipped with a Deltavision Core system, a 100× 1.4 NA Olympus lens, a Sedat Quad filter set (Semrock) and a CoolSnap HQ CCD Camera (Photometrics) controlled via softWoRx 4.1.0 software. Images were analyzed using ImageJ.

### Fluorescence in situ hybridization

Oligopaint FISH probe library MyTag was purchased from MYcroarray (see Supplementary Table 2). The library consisted of 477 high-stringency Atto-550 conjugated probes with an overall probe density of 13.9 probes per kb of the apicoplast genome. The probes were resuspended to 10 pmol/μl in ultrapure water (stock solution).

The fluorescence in situ hybridization protocol was adapted from (30). Parasites were washed twice with PBS and fixed with 10 volumes of the fixation solution (4% paraformaldehyde [Electron Microscopy Solutions #50-980-487], 0.08 % glutaraldehyde [Sigma #G6257] in PBS) for 1 h at 37° C. Fixed parasites were washed twice with PBS and permeabilized with 1 % Triton X-100 in PBS for 10 min at room temperature, followed by 3 washes in PBS. Next parasites were resuspended in the hybridization solution (50% v/v formamide [Sigma], 10% dextran sulfate [Millipore], 2× SSPE [Sigma], 250 mg/ml salmon sperm DNA [Sigma]) to approx. 20% hematocrit and incubated 30 min at 37° C. MyTag probes were resuspended in the hybridization solution to a final concentration of 1 pmol/μl, denatured for 5 min at 100° C and cooled on ice. 50 μl resuspended parasites were added to 20 μl hybridization solution with or without probes and incubated 30 min at 80° C followed by minimum 16 hours incubation at 37° C. Next parasites were subjected to following washes: 30 min at 37° C in 50 % (v/v) formamide, 2X SSC (Sigma); 10 min at 50° C in 1X SSC; 10 min at 50° C in 2X SSC; 10 min at 50° C in 4x SSC; 10 min at 50° C in PBS. Parasites were resuspended in 50 μl PBS, stained with 2 μg/ml Hoescht 33342 and imaged.

### Atg8 purification and anti-Atg8 antibody production

Hexahistidine-tagged Atg8 was expressed in Rosetta DE3 with rare codon plasmid from pRSF-1b-His-Atg8 (69, 70). Bacterial cultures were grown in the TB medium supplemented with 50 mg/l kanamycin and 34 mg/l chloramphenicol. At OD600=3 IPTG was added to the final concentration of 300 μM to induce Atg8 expression and cultures were further grown at 20° C for 16 hours. Bacteria were harvested by 20 min centrifugation at 800 g and bacterial pellets were resuspended in the buffer containing 50 mM HEPES pH 8.0, 500 mM NaCl, 1 mM MgCl2, 10 % glycerol and 2X Complete Protease Inhibitors (Pierce). Cells were lysed by a series of freeze-thaw cycles followed by passing them 3 times through the emulsifier Emulsiflex Avestin. Cell debris were removed by 30 min centrifugation at 30,000 × g. Clarified lysate was added to Talon resin (Clontech) and incubated 1 hr at 4° C. Beads were washed with the wash buffer containing 50 mM HEPES pH 8.0, 150 mM NaCl, 10 mM Imidazole pH 8.0 and 10 % glycerol. Protein was eluted with the wash buffer supplemented with 300 mM Imidazole pH 8.0, dialized against the wash buffer lacking imidazole, aliquoted and stored at −80 °C. Anti-Atg8 antibodies were raised in a rat and a guinea pig at Josman LLC. Josman is a licensed research facility through the USDA, number 93-R-0260 and has a PHS Assurance from the OLAW of the NIH, number A3404-01.

## Funding information

NIH 1K08AI097239 (Ellen Yeh)

Burroughs Wellcome Fund (Ellen Yeh)

NIH 1DP5OD012119 (Ellen Yeh)

Chan Zuckerberg Biohub (Ellen Yeh)

NIH 1DP2OD007124 (Jacquin C. Niles)

Bill and Melinda Gates Foundation (OPP1069759) (Jacquin C. Niles)

NIH P50 GM098792 (Jacquin C. Niles)

The funders had no role in study design, data collection and interpretation, or the decision to submit the work for publication.

## Acknowledgements

We thank Prof Karine LeRoch for the anti-Atg8 antibody which we used to validate anti-Atg8 antibodies produced in this study, Prof Walid Houry for the anti-ClpP antibody and Prof Aaron Straight for sharing the equipment. The pRSF-1b-His-Atg8 plasmid used to express His-Atg8 for immunization was a kind gift from Jürgen Bosch. We also thank Professors Pehr Harbury, Rajat Rohatgi, Lingyin Li, and Chao-Ting Wu for useful suggestions and comments on this project.This research was supported by grants from NIH and Burroughs Wellcome Fund to Ellen Yeh and from NIH and Bill and Melinda Gates Foundation to Jacquin Niles. Ellen Yeh is a Chan Zuckerberg Biohub investigator.

